# The MIF SNP rs755622 is a germline determinant of tumor immune activation in Glioblastoma

**DOI:** 10.1101/2022.03.07.483365

**Authors:** Tyler J. Alban, Matthew M. Grabowski, Balint Otvos, Defne Bayik, Ajay Zalavadia, Vlad Makarov, Katie Troike, Mary McGraw, Anja Rabljenovic, Adam Lauko, Chase Neumann, Gustavo Roversi, Kristin A. Waite, Gino Cioffi, Nirav Patil, Thuy T. Tran, Kathleen McCortney, Alicia Steffens, C. Marcela Diaz, J. Mark Brown, Kathleen M. Egan, Craig M. Horbinski, Jill S. Barnholtz-Sloan, Michael A. Vogelbaum, Richard Bucala, Timothy A. Chan, Manmeet S. Ahluwalia, Justin D. Lathia

## Abstract

While immunotherapies have shown durable responses for multiple tumors, their efficacy remains limited in some advanced cancers, including glioblastoma. This may be due to differences in the immune landscape, as the glioblastoma microenvironment strongly favors immunosuppressive myeloid cells, which are linked to an elevation in immune-suppressive cytokines, including macrophage migration inhibitory factor (MIF). We now find that a single-nucleotide polymorphism (SNP) rs755622 in the MIF promoter associates with increased leukocyte infiltration in glioblastoma. Furthermore, we identified lactotransferrin expression as being associated with the rs755622 SNP, which could also be used as a biomarker for immune infiltrated tumors. These findings provide the first example in glioblastoma of a germline SNP that underlies differences in the immune microenvironment and identifies high lactotransferrin as a potential factor promoting immune activation.

## Introduction

Immunotherapeutic strategies to stimulate anti-cancer immune responses have provided new treatment options in multiple advanced cancers(1–4). However, the efficacy of these approaches is variable, and in some tumors, such as glioblastoma (GBM), immunotherapy success has been limited(5–7). The obstacles to immunotherapy effectiveness in GBM include a highly suppressive myeloid cell-driven tumor microenvironment and systemic immune suppression, which limits T cell infiltration and activation, and the anatomical limitations of the blood-brain and blood-tumor barriers(8–12). Accordingly, identifying how resistance to these therapies is regulated is essential for developing effective next-generation immunotherapeutic strategies for GBM and other refractory cancers.

Within the GBM microenvironment, a series of cell-cell interactions concomitantly drive tumor growth and immune suppression(10). We previously identified an immune-suppressive pathway in GBM that is driven by macrophage migration inhibitory factor (MIF) secreted by cancer stem cells (CSCs) that in turn activates myeloid-derived suppressor cells (MDSCs)(13). Recent work from our laboratory and others has shown that MDSCs are increased in the circulation and tumor microenvironment(12), they portend a poor prognosis(8), their expansion can be driven by CSC-derived MIF(8, 13), and they can be reduced by MIF neutralization (either genetically or pharmacologically)(9, 14). Furthermore, MIF has been studied in a variety of cancers in the context of inflammation and has been found to regulate immune activity(15–32). However, MIF has not been explored in the context of immunotherapy.

MIF expression not only driven by disease state but also by functional germline genetic polymorphisms, A well-characterized MIF promoter SNP, rs755622, has been associated with multiple inflammatory conditions such as lupus, atopy, rheumatoid arthritis, septic shock, and cardiovascular disease^15–32^. This MIF SNP is in linkage disequilibrium with the presence of seven CATT repeats at the −794 promoter microsatellite, which leads to tighter binding by the transcription factor ICBP90 when compared to the more commonly occurring five or six CATT repeats(4). Accordingly, rs755622 is commonly analyzed in place of the −794 CATT microsatellite (27). While the minor allele frequency of the rs755622 MIF SNP ranges from 15-20% in Caucasians and >45% in individuals of African descent, it has not been associated with GBM growth or survival in large-scale genome-wide association studies(33). Given the role of MIF in immune response, we hypothesized that this MIF SNP may be associated with immunotherapy response in GBM. Here, we determined that patients with the MIF SNP rs755622 have an altered tumor microenvironment characterized by increased lymphocyte infiltration and enhanced lactotransferrin (LTF) expression.

## Results

### Patients with the MIF SNP rs755622 have an increase in Lactotransferrin (LTF) and increased immune microenvironment signaling

Based on our previous assessments of MIF as a driver of CSC-MDSC-mediated communication (13), we assessed overall MIF expression levels across brain tumors and found elevated MIF in isocitrate dehydrogenase (IDH) wild-type GBM patient tumor samples when compared to that of patients with lower-grade (IDH mutant astrocytomas and oligodendrogliomas) gliomas (**Fig. 1A**). Furthermore, the Genotype-Tissue Expression (GTEx) project identifies the MIF snp rs755622 as an expression quantitative trait loci highly associated with expression in most tissues except the central nervous system, where it has almost opposing results (**Supplemental Fig 1**). Using this information we hypothesized that GBM patients may have an increased prevalence of the regulatory MIF rs755622 nucleotide −173 G/C SNP (**Fig. 1B**). We assessed the rs755622 MIF SNP in three separate, annotated clinical cohorts of GBM patients (total of 966 patients including 449 from Cleveland Clinic, 386 from Moffitt Cancer Center, and 131 from Case Western Reserve University/University Hospitals of Cleveland). Our analysis of individual and combined cohort statistics revealed significant differences in the frequency of key prognostic indicators between cohorts (Karnofsky Performance Score (KPS), total surgical resection, receipt of standard of care (SOC), and recurrence) across the 3 cohorts. **Supplemental Fig. 2A-B**). We genotyped all patients and found a similar frequency of MIF SNP rs755622 major (G/G) and minor allele-containing (C/C or C/G, noted as C/^*^) genotypes across cohorts (**Supplemental Fig. 2C**) and when compared to the 1000 Genomes Project among those with European ancestry (**Supplemental Fig. 2C**). Additionally, we found strong linkage disequilibrium between the rs755622 SNP and the CATT repeats rs5844572 (**Supplemental Fig. 3A,B**) >90% cosegregation between CATT repeats 7-8 (rs5844572) and the MIF snp (rs755622), as previously reported(34). When evaluated according to major clinical prognostic indicators for GBM, we observed significant associations of sex and receipt of standard of care with the rs755622 MIF SNP genotype using descriptive analysis (**Supplemental Fig. 2B, D,E, Supplemental Fig. 4**). Univariate and multivariable survival analyses did not demonstrate any differences in overall or progression-free survival according to rs75622 MIF SNP genotype in any individual cohort or when data from all 3 cohorts were combined (**Supplemental Fig. 2D-F; Supplemental Fig. 4, 5**).

**Figure 1.**
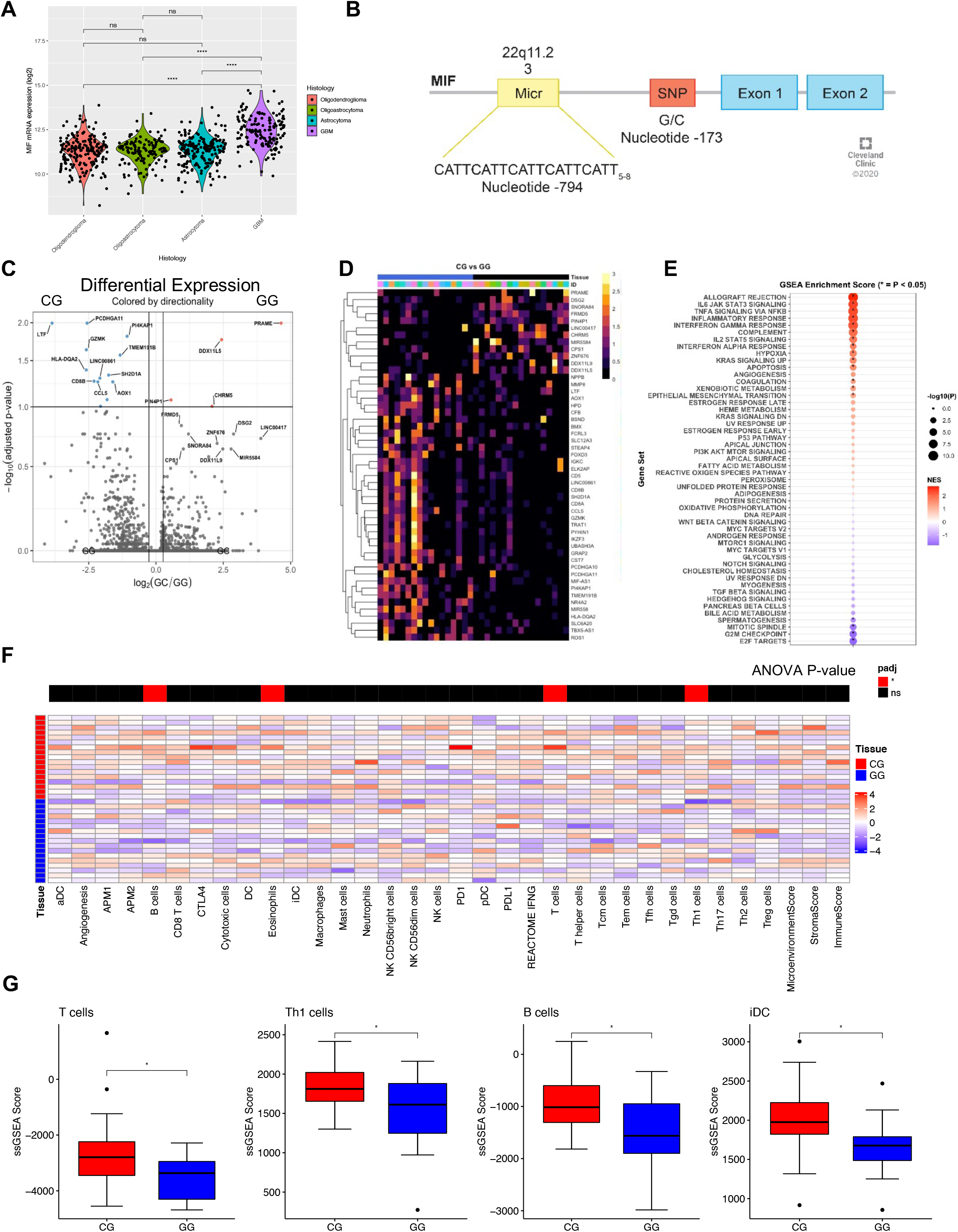
Patients with the MIF SNP rs755622 have an increase in Lactotransferrin (LTF) and increased immune microenvironment signaling. (**A**) Expression of the MIF gene mRNA in TCGA_GBMLGG was compared among the histology subtypes astrocytoma, oligodendroglioma, oligoastrocytoma, and GBM using unpaired t-test. (**B**) The MIF gene structure highlighting the −794 CATT repeat, which contains between 5 and 8 repeats, and the MIF SNP rs755622 at position −173. (**C**) Using GBM samples from n=17 G/G genotype patients and n=17 C/G patients, differential expression analysis was performed using DESeq2 on raw counts and the log fold change genes >1 and with p value >1 −log10(adjusted p-value). (**D**) The 25 genes with both the largest increases and largest decreases in gene expression by fold change are shown via heatmap, and rows are clustered using hierarchical clustering; transcripts per million are scaled to row for heatmap color scale. (**E**) GSEA (gene set enrichment analysis) was performed using Hallmark gene sets on the differentially expressed gene list comparing the C/G genotype to the G/G genotype sorted by log2 fold change for gene rank position. NES score is shown in red/blue with red enriched in C/G and blue enriched in G/G, while the size of the circle represents the −log10 p value as determined by GSEA. ssGSEA analysis was performed using the R package GSVA for cell type gene signatures to deconvolute immune cell types, and microenvironment, stromal, and immune scores were generated using xCell deconvolution R package from the bulk RNA-seq available for each of the n=34 patients. (**F**) ANOVA analysis was performed to compare the deconvolution scores between C/G and G/G patients, with the p value shown above the heatmap of the scaled scores (p>0.05=ns, p<0.05=*, p<0.01=**, p<0.001=***). (**G**) ssGSEA scores for significant populations are shown as a boxplot with unpaired t-test p values shown for each (p>0.05=ns, p<0.05=*, p<0.01=**, p<0.001=***, Two-Tailed T-Test).

Analyzing individual cohorts in univariate analyses and in combined analyses we observed no significant difference in GBM incidence or patient survival between the rs755622 MIF SNP genotypes, but we hypothesized that there may be differences in tumor and microenvironment interactions between genotypes given the association of the rs755622 MIF SNP with inflammation in non-oncologic conditions (**Supplemental Table 1**). To explore this possibility, we selected 17 patients with primary, untreated GBM from each MIF genotype in our Cleveland Clinic cohort with similar clinical parameters and outcomes (**Supplemental Table 2**) and subjected their tumor tissue to bulk RNA-sequencing. Differential gene expression analysis revealed that the rs755622 minor allele patients (*e.g*., −173 C SNP) had an enrichment in immune cell related genes (**Fig. 1C, D**). Gene set enrichment analysis (GSEA) utilizing the Hallmark curated gene sets identified a significant increase in inflammatory pathways in the minor allele patients (**Fig. 1E**). Additionally, ingenuity pathway analysis (IPA) showed similar findings, with increased innate immune response and prostaglandin signaling among the top enriched pathways of patients with the MIF SNP minor allele (**Supplemental Fig. 6**). In seeking to better understand the individual immune cell types that were changed between genotypes, we used a deconvolution approach using ssGSEA with curated gene sets and found a significant increase in cell signatures associated with an immune response in CG compared to GG patients as indicated by the anova p-value bar above each column of the heatmap (**Fig. 1F, 1G**).

Of the differentially expressed genes, lactotransferrin (LTF) was the most significantly upregulated in patients with the minor allele and positively correlated with increased T cell and immune responses (**Fig. 1C**). LTF is considered a key factor in first line immune defense against bacteria, yeast, viruses, fungi, and parasites, and may additionally contribute to the anti-tumor response(35–37). LTF is an acute phase immune mediator released from neutrophils and participates in the switch from innate to adaptive immune response. LTF signals through Toll-like receptors in myeloid cells to activate NFkB and CD40 expression and promote the initiation of anti-tumor immune responses(38, 39). It has also been associated with M1 polarization and in a pan-cancer study negatively correlated with tumor mutational burden(40). Taken together, these analyses suggest that while the rs755622 MIF SNP did not associate with differences in GBM incidence or survival, it did correlate with a difference in the immune cell composition within the tumor microenvironment.

### Immunofluorescence confirms enhanced T cell infiltration and CD8 T cell activation in GBM patients with the MIF SNP rs755622

To further interrogate the immune cell differences between MIF genotypes, we utilized matched tissue from 22 patients (11 C/^*^ and 11 G/G) from the same cohort subjected to RNA-sequencing. Staining for LTF confirmed the RNA-sequencing analysis and identified that LTF protein was increased in minor allele patients (**Fig. 2A-C**). The percent of LTF-expressing cells was further enhanced in minor allele patients with short term survival, based on an overall survival of less than the median of the 34 samples in this cohort (**Fig. 2D**). Additional assessment of T cell populations identified significant increases in CD8+ T cells in minor allele patients (**Fig. 2E-F**). Additionally, CD8+ T cells were less impacted by prognosis (**Fig. 2G**). While analysis of the activation marker CD107a on CD8+ T cells showed no difference between the good and poor prognosis groups (**Fig. 2H**), CD107a was significantly increased in CD8+ T cells in the minor allele patients (**Fig. 2I**). These data support the conclusion of an increase in immune infiltration of a cytotoxic T cell population in the minor allele patients along with enhanced LTF expression in the tumor microenvironment.

**Figure 2.**
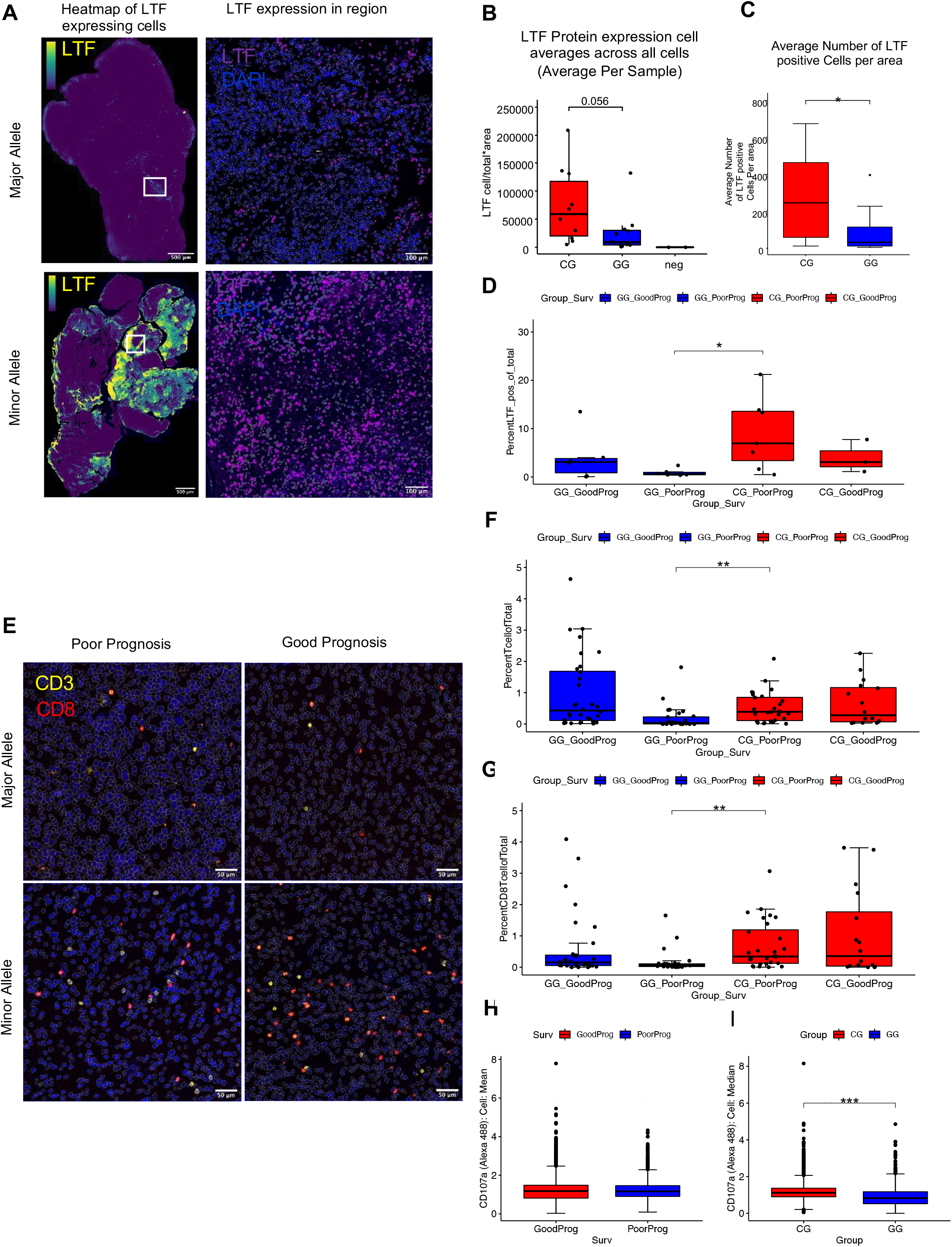
Immunofluorescence confirms enhanced T cell infiltration and CD8 T cell activation in GBM patients with the MIF SNP rs755622. (**A**) LTF expression was analyzed by immunofluorescence for n=22 samples matched from the RNAseq analysis to compare patients with the minor allele (C/G) and those with the major allele (G/G), with representative images of the dataset shown as a whole view of the slide in the heatmap images (left column, yellow=increased density of staining). In a 20x image of the slide, cells are outlined in white, with nuclei marked by DAPI and LTF pseudocolored purple. (**B**) The average LTF expression (mean fluorescence intensity) of cells per sample was compared between each genotype and the negative secondary-only control. (**C**) The quantity of LTF-positive cells per total area was measured and then compared between the C/G and G/G genotypes using unpaired t-test. (**D**) The percent of LTF-positive cells of the total cells per sample was compared between each genotype group and further subdivided into prognostic categories based on greater than or less than median overall survival. (**E**) Staining for CD3 and CD8 markers to determine CD8+ T cell infiltration, with representative images of each genotype and prognosis group (CD3=yellow, and CD8=red, DAPI=blue). Percent of T cells (**F**) and CD8+ T cells (**G**) of total cells per sample comparing the genotype/prognosis categories only. (**H**) CD107a expression of CD8+ T cells from all samples was compared between prognoses. (**I**) CD107a expression of CD8+ T cells was compared between the C/G genotype and G/G genotype. All statistics for this figures are performed using Two-tailed T-test (p>0.05=ns, p<0.05=*, p<0.01=**, p<0.001=***).

While we observed changes in the lymphoid compartment from the RNA-sequencing studies and validated these in human patient samples, these initial assessments did not focus on specific myeloid cell subtypes that are known to be involved in GBM immune suppression (including microglia, monocytes, macrophages, and MDSCs). Using matched samples from the RNA sequencing study, we next stained for CD4+ T cells and for myeloid cell subtypes known to be involved in GBM immune suppression (*e.g*., microglia, monocytes, macrophages, and MDSCs (n=11 C/* and n=11 G/G). Individual cell subtypes were identified using the top quartile for lineagespecific markers (CD4+ T cells: CD4^+^, CD11b^-^; macrophage: CD11b^+^, CD68^+^, HLA-DR+; microglia: P2RY12^+^; MDSCs: CD11b^+^, CD74^+^, CD68^-^, HLA-DR^-^; monocytes: CD11b^+^, HLA-DRA^+^, P2RY12^-^, CD68^-^) (**Fig. 3A, B**). Quantification revealed decreased macrophages in minor allele patients but no other major changes in MDSCs, ratio of MDSCs to CD8+ T cells, microglia, monocytes, and CD4+ T cells between genotypes (**Fig. 3C**). Due to the quantitative difficulties of assessing all myeloid cell lineages in an individual panel with lymphoid populations, we correlated cell types across samples per genotype (**Fig. 3D, E**). Using this approach, we found striking differences between genotypes, with the minor allele patients exhibiting an increase in CD8+ T cells and reductions in the myeloid compartment. Additionally, we found that LTF positively correlated with microglia and negatively correlated with the ratio of MDSCs to CD8+ T cells. These data lead to the overall conclusion that minor allele patients have increased lymphocyte infiltration with reduced macrophage content and further that LTF is associated with increased CD8+ T cells.

**Figure 3.**
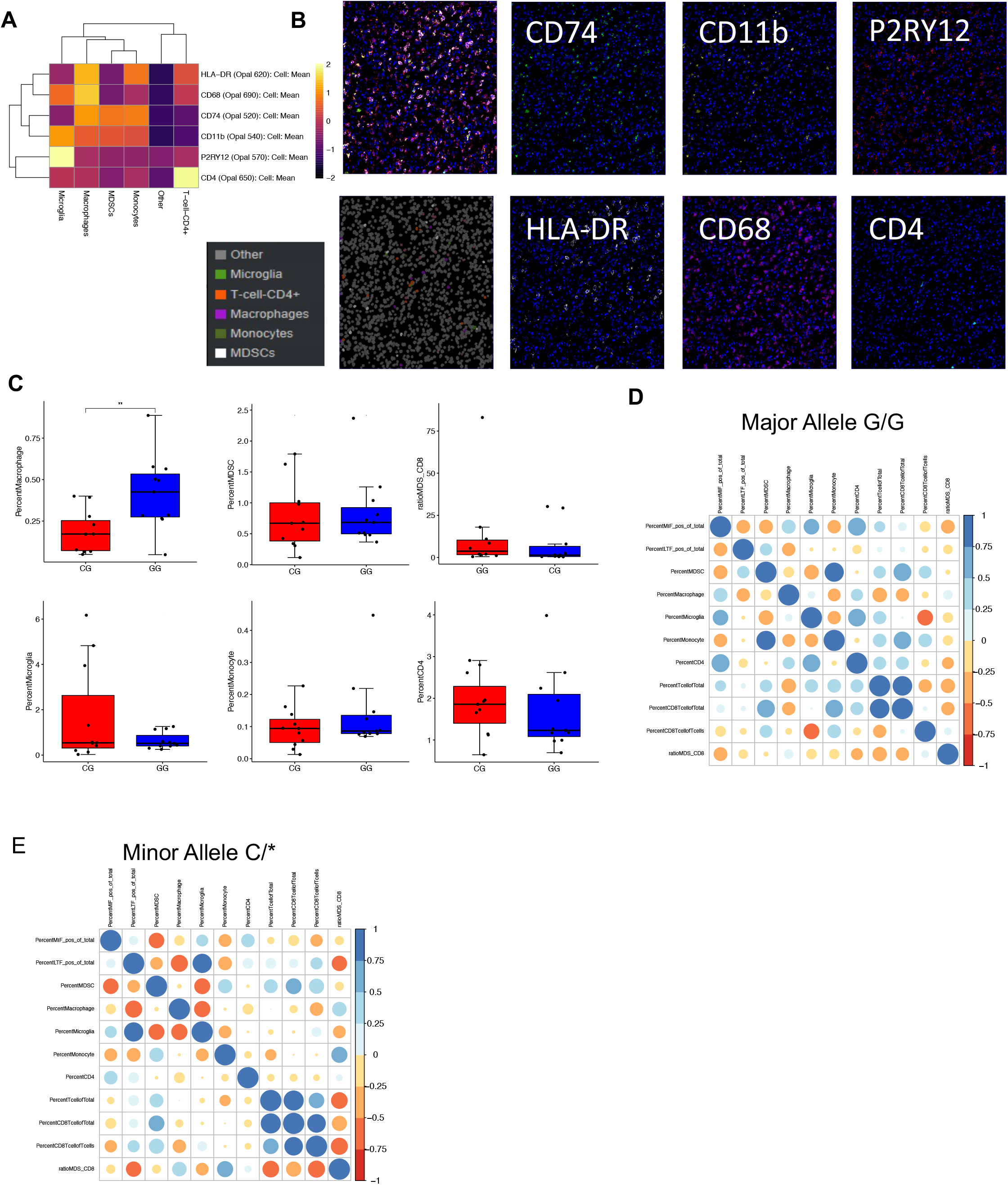
The MIF SNP rs755622 correlates with reduced macrophages and increased T cell infiltration by immunofluorescence. (**A**) A multiplex myeloid panel of antibodies was developed and included HLA-DR, CD68, CD74, CD11b, P2Ry12, and CD4, with the average expression shown for each cell type identified. (**B**) Representative images of each marker and pseudocoloring of each final cell type is shown in one representative image containing all cell types. (**C**) Analysis of the percent of macrophages, MDSCs, ratio of MDSC/CD8+ T cells, microglia, monocytes, and CD4+ T cells was performed using the percent of each cell type per sample out of total cells identified. (**D**) Correlation analyses were performed for each patient’s immune cell infiltrates from major allele genotype (G/G) patients using LTF staining, myeloid staining, and lymphocyte staining. Color scale and circle size are representative of the Pearson correlation coefficient. (**E**) Correlation analysis of staining cohorts for only the minor allele patients (C/*). All statistics for this figures are performed using Two-tailed T-test (p>0.05=ns, p<0.05=*, p<0.01=**, p<0.001=***).

### GBM patients with high LTF expression are immunologically activated

Seeking to expand on these initial observations, we sought to identify the rs755622 SNP in The Cancer Genome Atlas (TCGA) dataset but were unsuccessful because this marker is too far upstream of transcription to have read coverage in whole exome sequencing data. Given the strong correlation between the MIF minor allele with LTF expression, we used LTF expression as a surrogate of MIF genotype and interrogated immune changes associated with LTF. Notably, we found a similar differential expression profile between LTF-high (top 25%) and LTF-low (bottom 25%) patients in the TCGA as we did between genotypes in our dataset (**Fig. 4A**). In agreement with our initial assessment, GSEA revealed an increase in immune activation pathways in the LTF high expression patients, including allograft rejection and complement signaling, and a reduction in cancer-related pathways, including mitotic spindle and myc targets (**Fig. 4B, C**). To further identify cell type estimates between LTF-high and LTF-low samples, we performed deconvolution analysis and found increased immune and microenvironment scores, interferon gamma score, and macrophage content (**Fig. 4D, E**).

**Figure 4.**
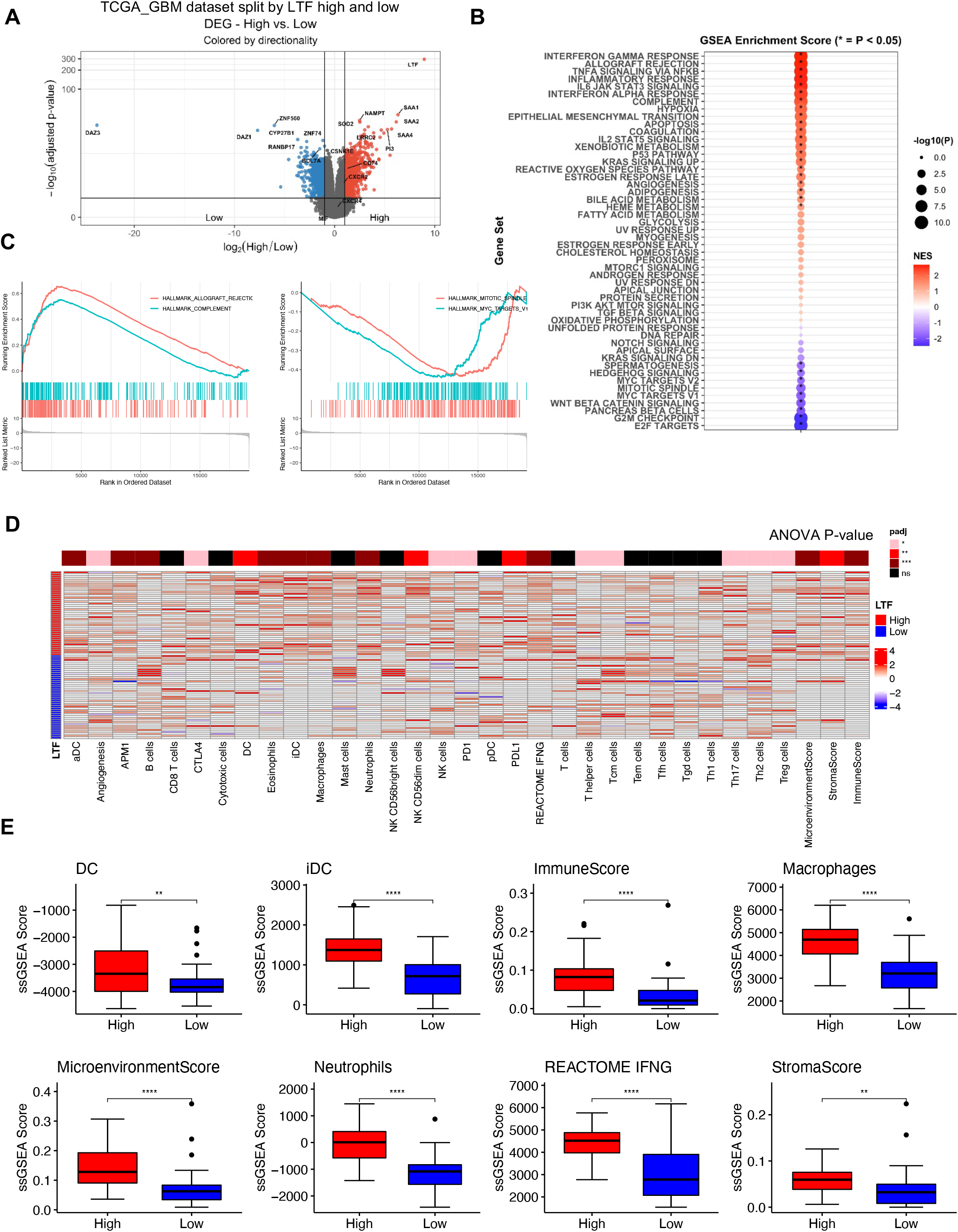
GBM patients with high LTF expression are immunologically activated. (**A**) TCGA_GBM mRNA seq data was analyzed to compare LTF-high (top 25% expression) and LTF-low (bottom 25% expression) patients, and differentially expressed genes are show between the groups. (**B**) GSEA analysis was performed based on the ranked gene list from differential expression between LTF-high and LTF-low samples, with pathways enriched in LTF-high samples shown in red. (**C**) Highlighted GSEA plots of rank ordered genes from the Hallmark pathways for allograph rejection, complement, which are enriched in LTF-high patients, and Mitotic Spindle and Myc Targets pathways enriched in the LTF-low patients. (**D**) Deconvolution analysis of the samples belonging the LTF-high and LTF-low groups showed increased immune cell type infiltration and increased immune scores, with ANOVA analysis for multiple comparisons showing the p-value by heatmap coloring. (**E**) Individual plots for significantly different estimated cell types and scores shown as individual boxplots with unpaired t-test.

### rs2096525 as a surrogate for rs755622 identifies increased immune activation in GBM

To determine whether these findings with respect to LTF signature recapitulate the rs755622 genotype, we examined possible associations with the rs2096525 SNP, which is in linkage disequilibrium with rs755622 but located within the first or second MIF intron. We interrogated the TCGA GBM whole exome sequencing dataset for the rs2096525 SNP and we found the minor allele to be present in approximately 12% of GBM patients (**Fig 5A**). Differential expression did not identify LTF, MIF or the MIF receptors as being differentially expressed (**Fig 5B**), however functional gene set enrichment analysis identified a similar increase in immune activation (**Fig 5C,D**). Additionally, a deconvolution analysis demonstrated similar increases in macrophages, neutrophils, and T cells for this SNP as observed for rs755622 RNA-seq study of CCF patients (**Fig 5E**). While the well characterized snp rs755622 has been demonstrated to play a major role in inflammatory conditions, here our data demonstrates that in glioblastoma those patients which have the minor allele have increased inflammation compared to the major allele counterparts.

**Figure 5.**
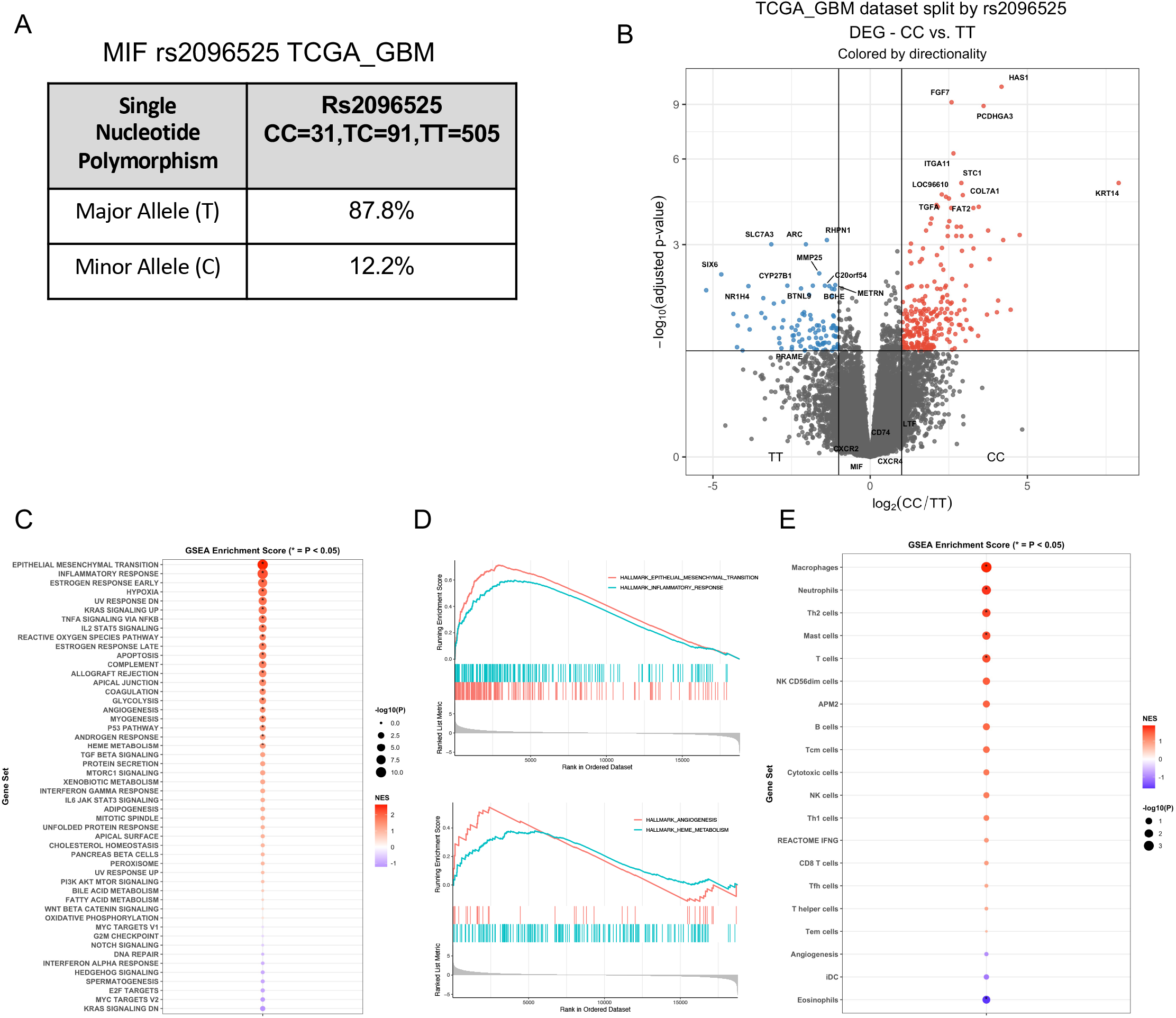
TCGA_GBM analysis of the MIF SNP rs2096525 identifies increased immune signaling. (**A**) Analysis of TCGA_GBM whole exome sequencing for rs2096525 identified 12.2% of patients with the minor allele. (**B**) The TCGA_GBM mRNA sequencing data and genotype information obtained from the whole exome sequencing data (n=159) were used for differential expression analysis between patients with the minor allele and patients homozygous for the major allele. Volcano plot represents deseq2 analysis of differential expression between the patients with the rs2096525 minor allele and patients homozygous for the major allele. (**C**) GSEA analysis of Hallmark pathways from differential expression analysis of rs2096525 patients demonstrates increased pathways in SNP-containing patients. (**D**) Selected enriched top two pathways are shown for GSEA analysis, with ranked genes in each pathway for enrichment and the top two downregulated pathways also shown. (**E**) Deconvolution analysis using ssGSEA with cell type gene sets from Nanostring demonstrates increased macrophages, neutrophils, and T cell populations in samples from patients with the minor allele rs2096525 SNP.

## Discussion

MIF, considered the first cytokine activity(41), has been extensively studied in the context of immune activation and the inflammatory response, as well as in tumor biology, where it has been shown to drive cancer cell proliferation and the generation of a tumor-promoting immune microenvironment(42). In GBM, MIF functions include enhancement of CSC maintenance(43, 44), resistance to therapies including standard-of-care therapy temozolomide(45) as well as the anti-angiogenic agent bevacizumab(46), as well as altering growth factor receptor signaling(38). However, the potential functional consequences of common MIF promoter polymorphisms, such as the −173 SNP (rs755622), have not been well studied in inflammation associated with malignancies. While the rs755622 SNP is associated with numerous inflammatory conditions and certain cancers, particularly those sensitive to immunotherapies(19, 42, 47, 48), we found that in GBM, there was no correlation with incidence or prognosis in response to standard-of-care therapy across three studied cross-institutional cohorts. This finding extended to the functional MIF promoter −794 CATT microsatellite (rs5844572) that is in linkage disequilibrium with the rs755622 SNP. We also observed no major difference in MIF level between genotypes, likely because MIF is already increased in GBM compared to normal, it is a secreted protein so RNA expression is not always accurate, IHC of the secreted MIF is difficult to accurately quantify, and MIF expression in GBM is further confound by many GBM patients being treated with dexamethasone, which increases MIF production(49). However, we found evidence for distinct tumor immune microenvironment between genotypes, with a heightened increase in CD8+ T cells in the minor allele patients. This enhancement in immune response parameters correlated with an enhancement in LTF expression in the minor allele patients.

Our initial assessment of MIF genotypes revealed an association between the minor SNP allele and LTF, which has not been previously described. In non-pathophysiological conditions, LTF is an iron binding glycoprotein that functions to protect against pathogens and has been shown to have anti-inflammatory activity. LTF has been described in cancer to function in an antiproliferative manner. In GBM, LTF expression is reduced compared to lower-grade brain tumors(50) and can inhibit GBM cell proliferation(51). LTF also has also been studied as a nanoparticle carrier for a variety of pre-clinical cancer therapies, including in GBM, where it has brain penetrance(52). In the context of MDSCs, we found that MIF enhances MDSC function in GBM and that LTF can induce MDSCs in pathological neonatal inflammatory conditions(53). While these two pathways have not been directly linked, there is a likely interaction based on our data. LTF and MIF are likely not co-regulated due to their location on separate chromosomes and their distinct f transcription factors, but future studies interrogating the molecular relationship between LTF and MIF may provide additional insight into the signaling networks that may functionally link these two proteins in GBM.

In analyzing the MIF snp rs2096525 which is in linkage disequilibrium with rs755622 we were initially surprised to not identify overlap of top differentially expressed genes with our RNAseq cohort. However, it is important to note that we utilized tru-seq methods of library preparation along with a much deeper sequencing strategy (targeted 100M total reads per sample). Additionally, TCGA-GBM samples have many batch effects such as institution, sequencers, and potential necrotic samples which are difficult to control for in a comparison like this. All of these factors combined make it challenging to interpret the direct overlap of top differentially expressed genes from TCGA with our cohort, but when using more broad approaches like gene set enrichment analysis we did see overlapping immune signatures similar to our cohort. This may indicate that overall, these samples are more immune activated, but mechanistic studies of LTF expression in relation to the MIF snp rs755622 and MIF itself are needed understand if LTF plays a direct role.

Utilizing both RNA-sequencing and matched tissue samples, we identified that MIF minor SNP allele patients who had increased LTF expression, also had an increase in CD8+ T cells and a reduction in macrophages, with no change in MDSCs. We also not observe a consistent change in tumor associated macrophages between MIF SNP patients and LTF expression, which could be due to a limitation of deconvolution methods in distinguishing myeloid subtypes. However, taken together, this immune microenvironment is likely more conductive to immune-activating strategies, and should be confirmed in future clinical trials. While our initial analysis revealed a high correlation between the MIF minor allele and elevated LTF, the association between the MIF SNP minor allele and LTF was made indirectly, due to the inability to efficiently identify the MIF SNP status in large genomic datasets based on its location in the promoter region, which is not covered by whole exome sequencing. Nonetheless, utilizing LTF at a median cut-off does not yield the same results as the top quartile of LTF, which more closely represents the frequency of the MIF SNP minor allele. Another limitation of our findings is that the MIF SNP has not been functionally characterized but is in linkage disequilibrium with the MIF −794 CATT repeat, which is associated with an increase in MIF production in immune cells and brain tissues.

The known genetic determinants of immunotherapy response in gliomas are currently limited to somatic mutations in IDH1/2, and the present findings identify a common germline SNP with clear immunologic significance that may be utilized to help improve clinical decision-making and support development of more effective immunotherapies.

## Materials and Methods

### Subjects and methods

For Cleveland Clinic, peripheral blood samples from 451 patients with GBM were collected through the Rose Ella Burkhardt Brain Tumor and Neuro-Oncology Center under approved IRB protocol 2559. White blood cells from each blood sample were isolated via Ficoll gradient and then snap frozen and stored at −80°C for research use. For this study, we selected all available GBM samples. For Moffitt Cancer Center, salivary DNA samples collected using Oragene kits was available for 386 recently diagnosed GBM patients under IRB protocol MCC 15004. DNA was extracted and stored in aliquot pellets at −80°C for future research use. For Case Western Reserve University/University Hospitals of Cleveland, peripheral blood samples from 131 patients with GBM were collected through the Ohio Brain Tumor Study at Case Western Reserve University, under approval from University Hospitals IRB CC296. Clinical and pathological data were gathered for each patient. Patient blood samples were collected and processed at the time of consent.

### DNA isolation and quantification

Genomic DNA was extracted from the peripheral blood of GBM patients using a Qiagen DNeasy Blood & Tissue Kit following the manufacturer’s protocols. DNA purity and concentration were measured using a ThermoFisher NanoDrop spectrophotometer.

### SNP genotyping

MIF SNP genotyping was performed using PCR amplification and subsequent restriction enzyme digestion with *AluI*. PCR was performed using Accuprime Pfx DNA polymerase (ThermoFisher, catalog number 12344-024) using 0.2 μmol forward primer (5’-CCCAAAGACAGGAGGTAC-3’) and 0.2 μmol reverse primer (5’-ATGATGGCAGAAGGACCAG-3’). PCR was run as follows: 95°C for 5 minutes, followed by 35 cycles of 95°C for 30 seconds, 60°C for 45 seconds, and 68°C for 1 minute. Following the 35 cycles, there was a final 68°C elongation step for 5 minutes, followed by storage at 4°C. After amplification, the PCR product was confirmed on a 1% agarose gel by identification of an approximately 500 bp product. After confirmation, 10 μl of PCR product was mixed with 2 μl 10x CutSmart buffer, 1 μl AluI, and 7 μl water and digested at 37°C for 1 hour. After digestion, the alleles containing a G (non SNP) produced a 450 bp fragment, while the alleles with a C (rs755622) produced a 270 bp fragment.

### SNP calling

Raw BAM files from the TCGA_GBM cohort were utilized to analyze the rs2096525 MIF SNP from whole exome sequencing data aligned by the TCGA. Aligned the SNP rs2096525 genotype was identified via use of HaplotypeCaller, where samples with alternative counts at the reference position chr22: 23894632 were identified. After classification of the samples by genotype, the phenotype data was downloaded via TCGA, and survival analysis was performed using log-rank test via R version 4.1.0.

### RNA sequencing

Flash-frozen tissue was requested from the Rose Ella Burkhardt Brain Tumor Center at the Cleveland Clinic under IRB 2559 and corresponded to n=34 patients with previously identified rs755622 SNP status from matched white blood cell pellets (n=17 C/*, and n=17 G/G). Within each group, patients were selected who underwent full Stupp protocol standard-of-care treatment and were evenly divided by sex and prognosis (<6 months progression-free survival (PFS) or >6 months PFS). Samples were processed and sequenced by Genewiz. Briefly, RNA was extracted by Qiagen RNeasy kit, and then the library was prepared using True-seq library preparation. Average sequencing depth was 40 Mbp per sample.

FASTQ files were aligned to the hg19 using STAR aligner with default parameters. Fragments were counted using Rsamtools with UCSC.hg19.knownGene transcript database. Raw counts were used in DESeq2 downstream for differential expression comparing the rs75662 SNP status groups.

### TCGA RNAseq data

Processed count data from TCGA_GBM mRNA dataset was downloaded from the Broad Firehose “http://gdac.broadinstitute.org/runs/stddata__2016_01_28/data/GBM/20160128/“, where the raw count file GBM.uncv2.mRNAseq_raw_counts.txt was utilized for downstream analysis.

### Differential expression

Raw counts from TCGA_GBM were analyzed using the R package DESeq2 version 1.29 in R version 4.0.1. After identification of the germline SNP status of rs2096525, the patient samples containing the minor allele were compared to patients homozygous for the major allele for differential expression.

### ssGSEA

The nCounter PanCancer Immune Profiling Panel gene set was used for immune infiltration deconvolution signatures. Each signature was used with ssGSEA using Gene Set Variation Analysis (GSVA) R package version 1.40.1. Microenvironment score, stromal score and immune score were generated using xCell R package version 1.1.0(54). Comparing signature scores between groups was performed using ANOVA with the p values shown for each comparison in the heatmaps above deconvolution heatmaps. All heatmaps of deconvolution and p-values were generated using pheatmap version 1.0.12.

### Immunofluorescence staining

Serial sections (7 μm thick) from each sample (formalin-fixed paraffin-embedded tumor biopsies) were stained with 3 different sets of markers and indicated below:

Set 1 (triple immunofluorescence staining): DAPI, CD3 (ab11089, abcam, 1:50), CD8 (85336, cell signaling, 1:50), CD107a (NBP2-52721, novus, 1:50)
Set 2 (double immunofluorescence staining): DAPI, MIF (MAB2892, R&D systems, 1:500), LTF (HPA059976, Atlas, 1:100)
Set 3 (multiplex staining): DAPI, CD74 (ab1794,Abcam/1:200), CD11b (ab133357,Abcam/1:200 P2RY12 (NBP2-33870,Novus Bio/1:200), HLA-DR (ab20181/Abcam/1:200), CD68 (790-2931, Ventana/1:200), CD4 (ab133616, Abcam/1:200)

For staining, slides were baked at 60^C^ prior to deparrifinization. Slides were then deparaffinized using a Leica Autostaine XL and antigen retrieval was performed using a sodium citrate buffer (pH6) with slides steamed in pressure cooker at 110^C^ for 10 minutes. Slides were then cooled to room temperature and transferred to water for 5 minutes prior to TBST for 15 minutes. Primary antibody placed at above concentrations and incubated in a humid chamber overnight at 4^C^. Slides placed on Biocare Intellipath Staining platform for blocking (3% donkey serum), secondary antibody incubations and Hoechst/DAPI staining (Thermo Scientific H1399/ 1mg/ml). The following secondary antibodies were used: Rabbit Cy3 (Jackson Immuno 711-165-152, 1:250), Mouse 488 (Jackson Immuno 715-545-151, 1:250), Rat Cy3 (Jackson Immuno 712-165-153, 1:250), Rabbit Cy5 (Jackson Immuno 711-175-152, 1:250). Slides manually cover slipped with an aqueous mounting medium.

### Imaging

Whole tissue sections were imaged with multispectral capabilities of the Vectra Polaris Automated Quantitative Pathology Imaging System (Akoya Bioscience Inc.). Multispectral images were then unmixed in inForm (Akoya Biosciences Inc., version 2.5) to obtain component images for each individual marker and tissue autofluorescence. Component image tiles were stitched and saved as OME-TIFF (Open Microscopy Environment) format for analysis, storage, and archival.

### Image analysis

The open source image analysis software QuPath was used for the detection and classification of cells. For each slide, 10 to 20 regions of interest (ROI) were selected to represent the different parts of the whole section while avoiding imaging, staining and sectioning artifacts. StarDist(55), a deep-learning algorithm, along with a pretrained model was used within QuPath for detecting cell nuclei from the DAPI channel. For each cell, intensity measurements were used to determine its positivity for each marker in the panel. A custom script with a manual decision tree(56) was implemented in QuPath to classify cells based on their positivity. Representative images were extracted using QuPath, and single-cell data for each sample were exported as .csv files for further analysis and charting in R version 4.0.1.

### Descriptive Statistics and Survival analysis

Demographic and clinical characteristics were evaluated between clinical cohorts. Analysis of variance (ANOVA) and chi squared tests were performed to assess differences in continuous and categorical variables, respectively. Additionally, these characteristics were assessed for MIF SNP rs755622. For this assessment, T tests were performed to assess differences in continuous data. All statistics were generated in R version 4.0.1.

### Survival analysis

Overall and progression-free survival of each clinical cohort was assessed for MIF snp rs755622. Kaplan Meier (KM) analysis was performed to evaluate the difference in survival and recurrence between GG genotype patients and CC or CG genotype patients. These analyses were also performed among only those cases who had received standard of care. Log rank tests were performed to assess differences in KM curves. Univariate and multivariable Cox proportional hazards models were generated to assess the impact of MIF snp rs755622 on overall and progression free survival. The proportional hazards assumption was assessed and not found in violation. Multivariable models were adjusted for age, sex, surgery, and standard of care. Hazard ratios (HR) and 95% confidence intervals (95% CI) are reported. All statistics were generated in R version 4.0.1.

## Author Contributions

**Conceived of the experiments:** T.J.A., M.M.G., B.O., M.A.V., R.B., T.A.C., M.S.A., J.D.L.; **Carried out experiments and analysis:** T.J.A., M.M.G., B.O., D.B., A.Z., V.M., K.T., M.M., A.R., A. L., C.N., G.R., K.A.W., G.C., N.P., T.T.T., K.M., A.S.; **Supervised and participated in analysis:** C.M.D., J.M.B., K.M.E., C.M.H. J.S. B.-S., M.A.V., R.B., T.A.C., M.S.A., J.D.L.; **Supervised the project and provided financial support:** M.A.V., R.B., T.A.C., M.S.A., J.D.L.; **Wrote the manuscript:** T.J.A., R.B., T.A.C., M.S.A., J.D.L.; **Final approval of manuscript:** All authors

## Acknowledgements

We thank Dr. Erin Mulkearns-Hubert for editorial assistance and members of the Lathia laboratory for insightful discussions. We thank also Ms. Amanda Mendelsohn for illustration assistance. This work was supported by National Institutes of Health grants R01NS109742 (J.D.L., M.A.V., R.B.), R35 CA232097 (T.A.C.), K99CA248611 (D.B.), F31 NS101771 (T.J.A.), and T32 CA059366 (T.J.A.), Cleveland Clinic Research Program Committees (B.O.), NIH K12CA215110 (T.T.) as well as the Lerner Research Institute (T.A.C., J.D.L.) and Case Comprehensive Cancer Center (T.A.C, J.M.B, T.A.C., J.D.L.).

## Supplemental Figure Legends

**Supplemental Figure 1.**
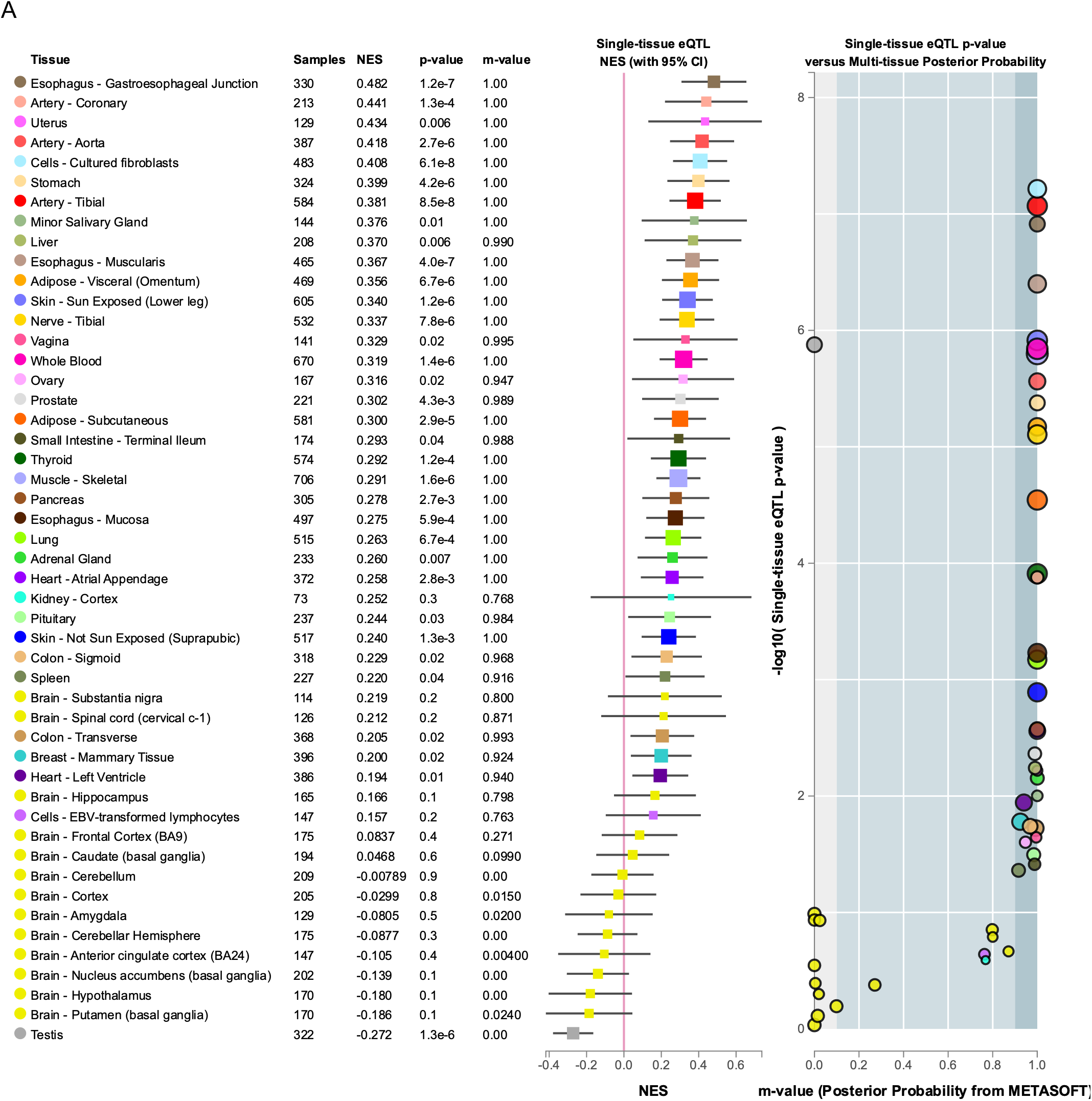
GTEX database analysis of MIF and rs755622 as expression quantitative loci across tissues. Searching the GTEX database for rs755622 and its association across multiple tissues for a link with MIF expression shows significance in many tissues except central nervous system related tissues.

**Supplemental Figure 2.**
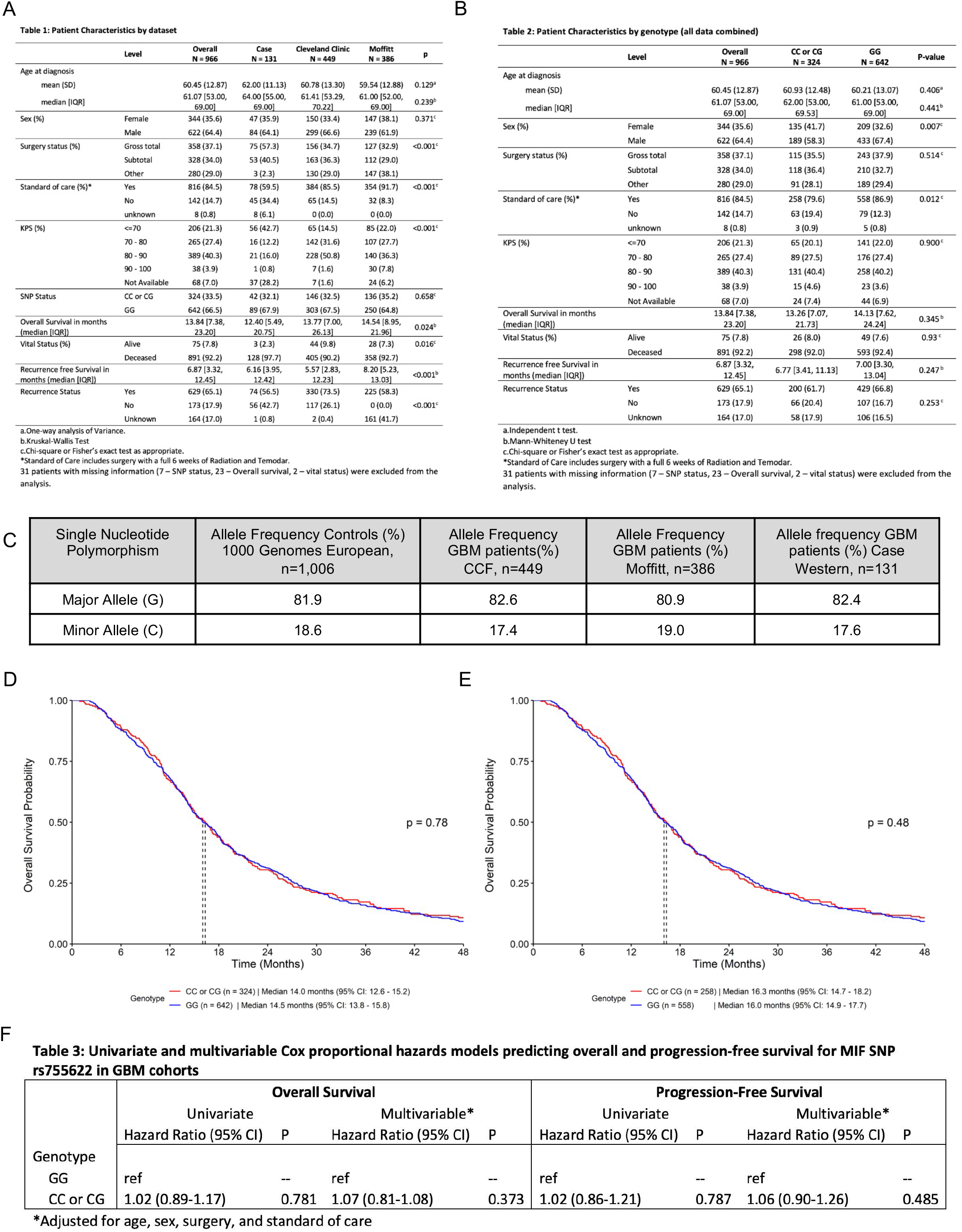
Univariate and multivariable analysis of the MIF SNP rs755622 in GBM cohorts. Germline DNA was acquired from PBMCs or saliva samples from n=449 (Cleveland Clinic), n=386 (Moffitt Cancer Center), and n=131 (Case Western) GBM patients and then tested for the MIF SNP rs755622 using PCR with restriction enzyme digestion as described in detail in the methods. (**A**) Descriptive statistics of clinical features of GBM including sex, surgery type, standard of care, KPS, and SNP status including all cohorts of GBM patients. Primary features known to associate with outcome such as higher KPS, total as compared to subtotal tumor resection and treatment with standard of care all confer a survival advantage. (**B**) Descriptive statistics combining data from all three cohorts of GBM patients demonstrated statistically significant differences in allele frequencies of the MIF SNP rs755622 (minor ‘C’ allele containing genotypes vs the homozygous major ‘G’ allele genotype) for standard of care treatment and sex status. (**C**) Allele frequencies in MIF rs755622 are similar in the 3 GBM cohorts and the 1000 genomes European cohort using Hardy-Weinberg principle and chi-square test for differences from reference control cohort. (**D**) Univariate analysis of overall survival across all cohorts shows no survival difference by log rank test. (**E**). Restricting to patients uniformly treated with standard of care, we observed no significant difference in overall survival according to MIF rs755622 genotype (CC/CG versus GG) by the log rank test. (**F**). Univariate and multivariable Cox proportional hazards models predicting overall and progression-free survival for MIF SNP rs755622 in GBM cohorts.

**Supplemental Figure 3.**
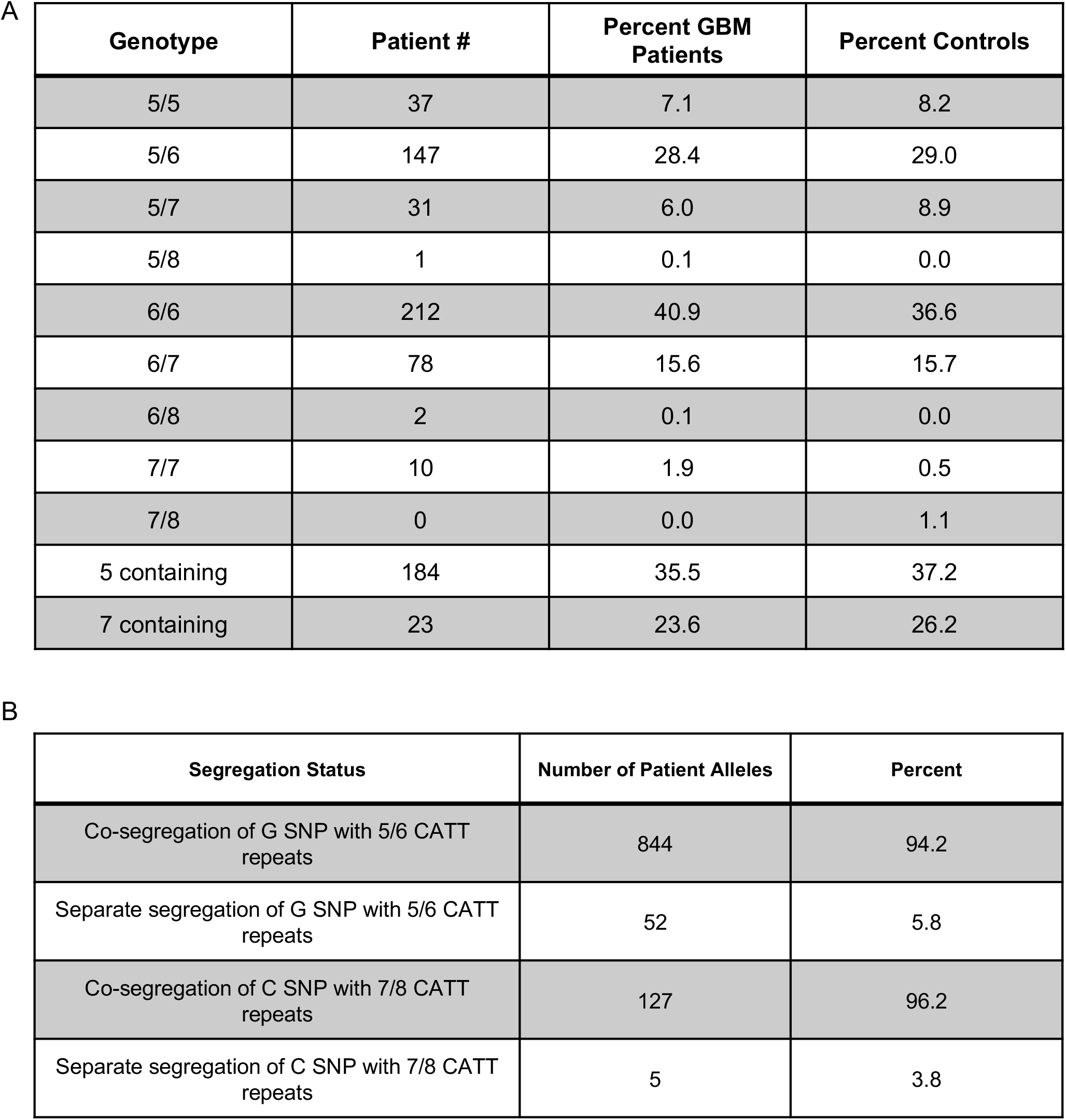
MIF CATT repeat analysis in GBM patients shows correlation with rs755622. (**A**) The CATT microsatellite in the MIF promoter upstream of the MIF SNP rs755622 was analyzed by capillary electrophoresis in the Cleveland Clinic cohort. (**B**) Those patients with both rs755622 SNP information and CATT repeat information were correlated to confirm linkage disequilibrium, which showed approximately 95% segregation of the 5/6 repeat and the major allele at rs755622, while 7/8 CATT repeats were linked with the minor allele of rs755622.

**Supplemental Figure 4.**
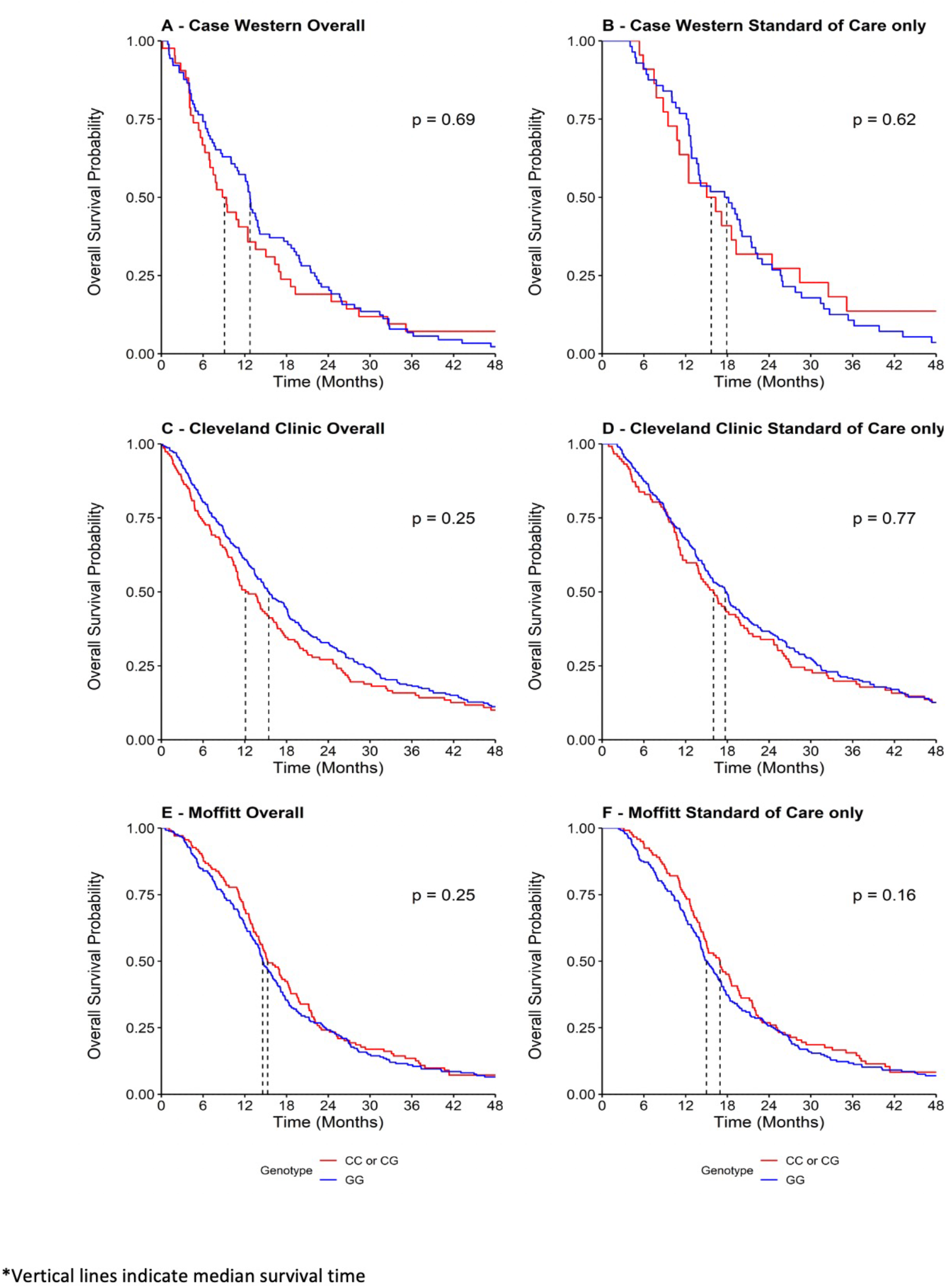
Univariate analysis of clinical cohorts for MIF snp rs755622 (Overall Survival). (**A-F**) Univariate analysis of overall survival comparing patients with the minor allele to patients homozygous for the major allele across all samples (left column) and then those who received full standard of care treatment protocol (right column), with log rank p-value comparing each group provided to the right of each curve.

**Supplemental Figure 5.**
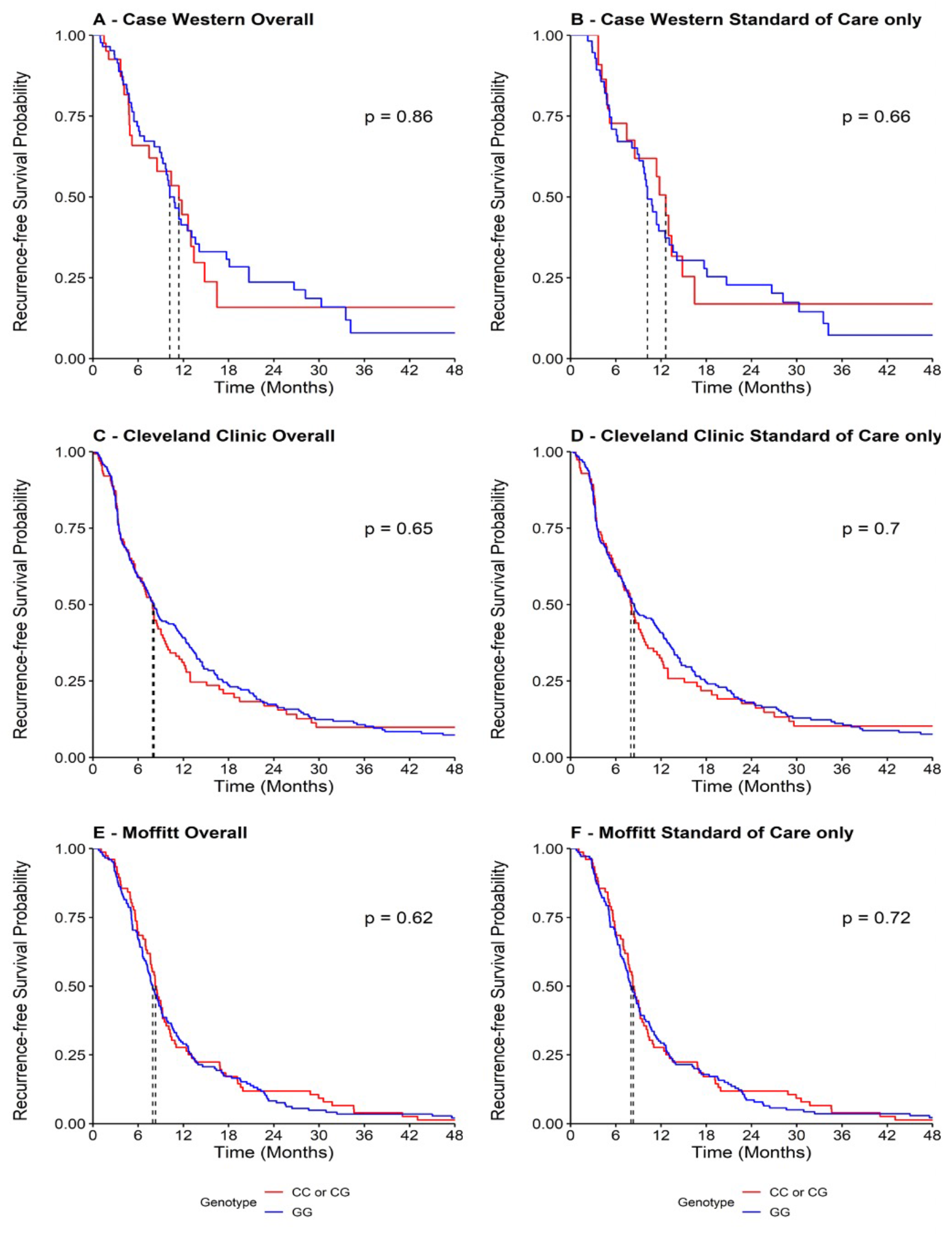
Univariate analysis of clinical cohorts for MIF snp rs755622 (Progression Free Survival). (**A-F**) Univariate analysis of recurrence-free survival comparing patients with the minor allele to patients homozygous for the major allele across all samples (left column) and then those who received full standard of care treatment protocol (right column), with log rank p value comparing each group provided to the right of each curve.

**Supplemental Figure 6.**
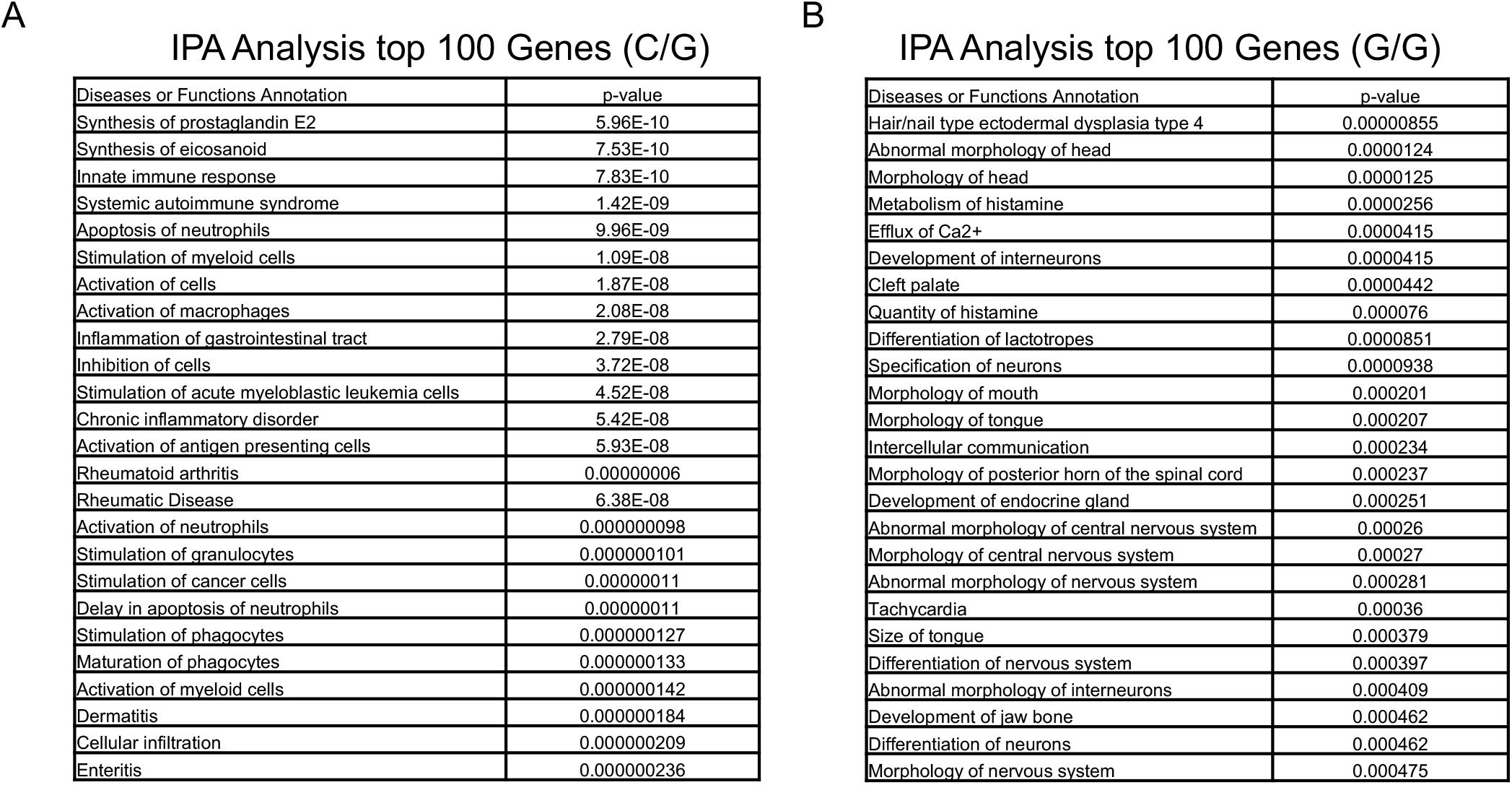
Ingenuity pathway analysis of C/G (rs755622) vs homozygous dominant. Genes from the differential expression analysis of rs755622 minor allele patients vs major allele patients were ranked by log fold change and the top100 genes increased in minor allele and top 100 decreased in minor allele were utilized for Ingenuity pathway analysis. (**A-B**) Pathway description and p-value are shown for the top 25 pathways associated with the 100 genes.

**Supplemental Table 1.** MIF SNP rs755622 highlighted literature in inflammatory associated conditions. Summary highlighting selected articles pertaining to rs755622 history and research in inflammatory conditions, and provides relative clinical importance of this germline SNP.

| PMID | Title |
| --- | --- |
| PMID: 19941661 | A MIF haplotype is associated with the outcome of patients with severe sepsis: a case control study |
| PMID: 20388640 | Predictors of response to intra-articular steroid injection in psoriatic arthritis |
| PMID: 20447688 | The MIF -173G/C polymorphism and risk of childhood acute lymphoblastic leukemia in a Chinese population |
| PMID: 16380915 | Replication of putative candidate-gene associations with rheumatoid arthritis in >4,000 samples from North America and Sweden: association of susceptibility with PTPN22, CTLA4, and PADI4 |
| PMID: 17585860 | Macrophage migration inhibitory factor in acute lung injury: expression, biomarker, and associations. |
| PMID: 20811626 | Genetic variants in inflammation-related genes are associated with radiation-induced toxicity following treatment for non-small cell lung cancer. |
| PMID: 23402792 | Macrophage migration inhibitory factor (MIF): genetic evidence for participation in early onset and early stage rheumatoid arthritis |
| PMID: 24151396 | Macrophage migration inhibitory factor gene polymorphisms in inflammatory bowel disease: An association study in New Zealand Caucasians and meta-analysis |
| PMID: 26107210 | Contribution of Macrophage Migration Inhibitory Factor -173G/C Gene Polymorphism to the Risk of Cancer in Chinese Population |
| PMID: 26542751 | Functional polymorphisms in the gene encoding macrophage migration inhibitory factor (MIF) are associated with active pulmonary tuberculosis |
| PMID: 29545822 | Macrophage Migration Inhibitory Factor -173 G/C Polymorphism: A Global Meta-Analysis across the Disease Spectrum. |
| PMID: 32560699 | MIF -173G/C (rs755622) polymorphism modulates coronary artery disease risk: evidence from a systematic meta-analysis. |
| PMID: 22113576 | Polymorphisms in immune function genes and non-Hodgkin lymphoma survival. |
| PMID: 24530749 | Macrophage migration inhibitory factor: association of -794 CATT5-8 and -173 G>C polymorphisms with TNF-α in systemic lupus erythematosus |
| PMID: 27696094 | A Macrophage Migration Inhibitory Factor Polymorphism Is Associated with Autoimmune Hepatitis Severity in US and Japanese Patients |
| PMID: 29661540 | MIF functional polymorphisms (-794 CATT 5-8 and -173 G>C) are associated with MIF serum levels, severity and progression in male multiple sclerosis from western Mexican population |
| PMID: 29996006 | MIF-173G/C (rs755622) polymorphism as a risk factor for acute lymphoblastic leukemia development in children |
| PMID: 30747392 | Macrophage migration inhibitory factor polymorphisms are a potential susceptibility marker in systemic sclerosis from southern Mexican population: association with MIF mRNA expression and cytokine profile |
| PMID: 31978276 | Association of the genetic variants (-794 CATT5-8 and -173 G > C) of macrophage migration inhibitory factor (MIF) with higher soluble levels of MIF and TNFα in women with breast cancer |

**Supplemental Table 2.** Clinical Cohort summary for samples used in RNA-sequencing study. Summary highlighting patients used for RNAseq analysis from ccf cohort n=17 G/* and n=17 G/G patients. Molecular markers and clinical data available were selected along with overall survival and sex to evenly distribute similar cohorts of patients and minimize confounders.

| Characteristic |  | CC, N = 1<sup>1</sup> | CG, N = 16<sup>1</sup> | GG, N = 17<sup>1</sup> |
| --- | --- | --- | --- | --- |
| race |  |  |  |  |
| black |  | 0 (0%) | 1 (6.2%) | 1 (5.9%) |
| non hispanic white |  | 1 (100%) | 15 (94%) | 16 (94%) |
| age |  | 59 (59, 59) | 64 (53, 69) | 59 (49, 66) |
| sex |  | 1 (100%) | 6 (38%) | 6 (35%) |
| kps.at.diagnosis | 60 | 0 (0%) | 1 (6.2%) | 0 (0%) |
|  | 70 | 0 (0%) | 1 (6.2%) | 1 (5.9%) |
|  | 80 | 0 (0%) | 8 (50%) | 4 (24%) |
|  | 90 | 1 (100%) | 6 (38%) | 12 (71%) |
| deletion_1p |  | 0 (0%) | 1 (6.2%) | 2 (12%) |
| deletion_19q |  | 0 (0%) | 5 (31%) | 1 (6.2%) |
| idh1_mutated |  | 0 (0%) | 0 (0%) | 1 (5.9%) |
| mgmt_methylated |  | 0 (NA%) | 2 (33%) | 4 (80%) |
| ki_67 |  | 5 (5, 5) | 25 (19, 32) | 25 (14, 32) |
| egfr_amplified |  | 0 (0%) | 9 (56%) | 5 (31%) |
| pfs |  | 19 (19, 19) | 8 (5, 21) | 25 (5, 51) |
| os |  | 27 (27, 27) | 12 (10, 25) | 36 (9, 82) |
| <sup>1</sup>n (%); Median (IQR) |  |  |  |  |

